# Seasonal and daily patterns in known dissolved metabolites in the northwestern Sargasso Sea

**DOI:** 10.1101/2022.12.21.521480

**Authors:** Krista Longnecker, Melissa C. Kido Soule, Gretchen J. Swarr, Rachel Parsons, Shuting Liu, Winifred M. Johnson, Brittany Widner, Ruth Curry, Craig A. Carlson, Elizabeth B. Kujawinski

## Abstract

Organic carbon in seawater plays a significant role in the global carbon cycle. The concentration and composition of dissolved organic carbon, operationally defined in this project as organic carbon that passes through a 0.2 µm filter, reflect the activity of the biological community and chemical reactions that occur in seawater. From 2016 to 2019, we repeatedly sampled the oligotrophic northwest Sargasso Sea in the vicinity of the Bermuda Atlantic Time-series Study site (BATS) to quantitatively follow known compounds within the pool of dissolved organic matter in the upper 1000 meters of the water column. Dissolved metabolite concentrations revealed patterns with depth and time with most metabolites showing surface enrichment and lower concentrations with increasing depth. Select metabolites displayed seasonal patterns throughout the year, and this seasonality was repeated in each of the years sampled. Concentrations of vitamins, including pantothenic acid (vitamin B_5_) and riboflavin (vitamin B_2_), increased annually during winter periods when mixed layer depths were deepest. During diel sampling, light-sensitive riboflavin decreased significantly during daylight hours. Metabolite concentrations over time at BATS showed less variability compared to a previous sample set collected over a broad latitudinal range in the western Atlantic Ocean. The metabolites examined in this study are all components of central carbon metabolism. By examining these metabolites at finer resolution and in a relatively long time series, we have insights into microbial biogeochemical activity, data which are fundamental to understanding the chemical response of marine systems to future changes in climate.

## Introduction

Dissolved organic carbon (DOC) in seawater is a significant fraction of carbon on Earth, with a concentration equivalent to the amount of carbon dioxide stored in the atmosphere (Siegenthaler and Sarmiento 1993). In marine systems, the amount of carbon present in dissolved form is 200 times the amount found in particulate form (Hansell et al. 2009). Over the last thirty years, scientists have made advances in quantifying where dissolved organic carbon is produced, consumed, and stored in the global ocean (Carlson and Hansell 2015). Microorganisms are the main consumers of DOC, and multiple researchers have examined the connections between the diversity of microorganisms in a marine system and their ability to consume dissolved organic matter (Bercovici et al. 2021; Liu et al. 2020a; Stephens et al. 2020). These and other interactions between biology and chemistry play a key role in defining the factors controlling organic carbon distributions in marine systems, both now and in the future ocean.

Time-series studies in the northwest Sargasso Sea extend back to 1954 at Hydrostation S and began in 1988 at the Bermuda Atlantic Time-series Study site (BATS). Repeated sampling at these sites has revealed long-term (multi-year) changes in properties such as temperature, dissolved inorganic carbon, and dissolved oxygen (Bates and Johnson 2020), and changes in the water masses in the vicinity of BATS (Stevens et al. 2020). These studies have further identified seasonally varying physical and biogeochemical processes (Bates et al. 1996; Michaels et al. 1994; Steinberg et al. 2001) that drive changes in organic carbon concentrations (Carlson et al. 1994; Hansell and Carlson 2001), nutrient cycling and carbon export (Lomas et al. 2013), carbon isotopes (Gruber et al. 1998), oxygen levels (Billheimer et al. 2021; Fawcett et al. 2018), cellular carbon quotas (Casey et al. 2013), and carbon export by phytoplankton (De Martini et al. 2018; Lomas et al. 2022). Seasonal variability at BATS is also evident in the biological community, including viruses (Parsons et al. 2012), autotrophic microorganisms (DuRand et al. 2001; Lomas et al. 2010), heterotrophic microorganisms (Morris et al. 2005; Treusch et al. 2009), and mesozooplankton (Blanco-Bercial 2020). Logically, temporal variability in these chemical and biological parameters should also be expressed in the distribution, quantity and composition of organic compounds found in the northwest Sargasso Sea.

Mass spectrometry has been used to identify the composition and patterns in the distribution of dissolved organic matter (Steen et al. 2020). Analytical advances, including pre-separation techniques such as liquid chromatography, have enabled both targeted investigations of select known organic compounds (Durham et al. 2015; Widner et al. 2021) and untargeted investigations that characterize previously unknown dissolved organic compounds in marine systems (Longnecker and Kujawinski 2017; Petras et al. 2021). Sinking particles contain compositionally distinct organic compounds compared to suspended particles (Johnson et al. 2020) and particulate metabolites can vary on diel timescales (Boysen et al. 2021). Furthermore, previous work in the Atlantic Ocean found that the composition of metabolites in suspended particulate organic material is distinct from the dissolved metabolites (Johnson et al. accepted, 2022). However, knowledge of the compounds that comprise dissolved organic matter in seawater has lagged studies of particulate organic matter partially due to the analytical challenges associated with concentrating, isolating, and analyzing dissolved metabolites. The research projects detailed above have provided important spatial details on the complex nature of dissolved organic matter in marine ecosystems but lack information on temporal variability.

As a component of BIOS-SCOPE, a multi-year, trans-disciplinary program to study microbial processes, structure and function in the Sargasso Sea, we have used targeted metabolomics to track and quantify a set of compounds central to cellular carbon metabolism over multiple time scales (diel to seasonal). These compounds were measured in the dissolved organic fraction collected over the upper 1000 meters at regular temporal intervals (monthly to seasonal) between 2016 and 2019. Each July, multi-day BIOS-SCOPE process cruises enabled higher frequency (6-hour) sampling to investigate diel patterns in the summer. The metabolite data have revealed recurring patterns in metabolite concentrations over depth and through time on both diel and seasonal timescales. The sampling location, near the BATS site, permits us to place this newly-generated metabolite data within the context of the rich monthly biogeochemical data collected at BATS since 1988. The present baseline study thus establishes the groundwork for using these compounds to track future changes in environmental conditions in oligotrophic oceans.

## Materials and Methods

### Hydrographic data

From July 2016 through July 2019, the upper ocean was sampled at three locations in the Sargasso Sea: BATS (31°40’N, 64°10’W), Hydrostation S (32°10’N, 64°30’W), and east of BATS (AE1916, 32°10’N, 64°13’W). On a subset of BATS cruises, water samples were collected from a minimum of six vertical levels spanning the surface down to 1000 m. During select times (July each year, September 2016, April 2017, and May 2019), the station was occupied for multiple days allowing for higher sampling frequency (Supplemental Table 1). Because local water mass structure responds to winter mixing, thermal stratification, mesoscale eddies and varying light penetration, we assigned sample depths to a vertical zonation of the water column (Curry et al. unpublished). We have samples from four seasons (mixed, spring, stratified, fall) with season boundaries set by the relative positions of the chlorophyll maximum layer and the mixed layer depth which was defined by a density threshold criteria as the depth where sigma-theta (σ_0_) exceeds the surface density by 0.125 kg m^-3^. We have samples from nine of the vertical zones defined by Curry et al., and we grouped our samples into the photic zone (VZ_0_, VZ_1_, and VZ_2_), the subphotic region (VZ_3_), and the deep ocean (VZ_4_ through VZ_10_). The details on the bounds used to define the vertical zones are detailed in Supplemental Table 3.

Water samples were collected using 12 L Ocean Test Equipment bottles mounted on a rosette equipped with a Conductivity-Temperature-Depth (CTD), fluorometer, and an oxygen sensor. Water samples were processed to obtain concentrations of particulate organic carbon, dissolved/total organic carbon, nutrients (nitrate + nitrite, phosphate), bacterial abundance using epifluorescence microscopy, and bacterial production via ^3^H-leucine incorporation (process cruises only) using established methods described in Knap et al. (1996), Halewood et al. (2022), and Liu et al. (2022). Both dissolved organic carbon (DOC) and total organic carbon (TOC) samples were collected during this project and the two are statistically indistinguishable in oligotrophic waters (Halewood et al. 2022); in this paper we refer to DOC or TOC concentrations as ‘bulk organic carbon’ concentrations.

### Metabolite extractions

Water (4 – 10 L) was collected directly from the sampling bottles into polytetrafluoroethylene (PTFE) or polycarbonate containers. Water was then filtered using a peristaltic pump through a 0.2 µm filter (Omnipore, EMD Millipore) held in a Teflon filter holder. The filtrate was acidified with 12 M HCl (to ∼pH 2-3). Dissolved organic molecules were extracted from the filtrate using solid phase extraction (SPE) with Agilent Bond Elut PPL cartridges (Dittmar et al. 2008; Longnecker et al. 2015). The cartridges were pre-conditioned with methanol and the filtrate was pulled through the cartridges via PTFE tubing using a vacuum pump. The cartridges were rinsed with ∼24 mL of 0.01 M HCl and then allowed to dry by pulling air over the cartridge for 5 min. The samples were eluted with methanol into a glass test tube. Extracts were evaporated to near dryness using a Vacufuge (Eppendorf) and reconstituted in 250 µL of 95:5 (v/v) water/acetonitrile or 100% water with isotopically-labeled compounds that serve as injection standards (Kido Soule et al. 2015).

### Targeted mass spectrometry

Samples were analyzed using ultra high performance liquid chromatography (Accela Open Autosampler and Accela 1250 Pump, Thermo Scientific) coupled to a heated electrospray ionization source (H-ESI) and a triple quadrupole mass spectrometer (TSQ Vantage, Thermo Scientific) operated under selected reaction monitoring (SRM) mode (Kido Soule et al. 2015). Separation was performed at 40 °C on a reversed phase column (Waters Aquity HSS T3, 2.1 x 100 mm, 1.8 μm) equipped with a Vanguard (Waters) pre-column. Mobile phase A was 0.1% formic acid in water and mobile phase B was 0.1% formic acid in acetonitrile. The flow rate was maintained at 0.5 mL min^-1^. The gradient began at 1% B for 1 minute, increased to 15% from 1 to 3 minutes, then increased to 50% B from 3 to 6 minutes, and then increased to 95% B from 6 to 9 minutes. The mobile phase was maintained at 95% B until 10 minutes and then decreased to 1% B from 10 to 10.2 minutes and held at 1% B for the remainder (12 minute total run time). Samples were run in both positive and negative ion modes using a 5 μL injection for each. We used two precursor-to-product ion transitions to identify each metabolite, one for quantification and a second to confirm metabolite identity. The complete list of metabolites analyzed in this project is provided in Supplemental Table 2.

Raw data files were converted to mzML format using msConvert (Chambers et al. 2012), and El-MAVEN (Agrawal et al. 2019) was used to identify and integrate peaks for all samples and standards. Metabolite concentrations were calculated using a standard curve of at least five points. For metabolites with an extraction efficiency greater than 1%, we corrected the measured concentrations using the data available from Johnson et al. (2017) and Swarr et al. (unpublished data) for metabolites added since 2017. Quality control checks for peak shape and instrument response were implemented as described in Kido Soule et al. (2015).

### Pre-extraction derivatization to enhance capture of polar dissolved metabolites

In May 2019 we had an opportunity to use a new method developed by Widner et al. (2021) that derivatizes metabolites in filtered seawater prior to solid phase extraction to improve their extraction efficiency and quantification. We processed a subset of samples by this method, replicate samples from 8 depths (cast 6) and 12 depths (cast 8). Briefly, sodium hydroxide (8M) was added to filtered seawater, followed by the addition of 5% v/v benzoyl chloride in acetone. After shaking, samples were neutralized with concentrated phosphoric acid and stored frozen until processing on land through solid phase extraction to remove salts. The samples were then analyzed by liquid chromatography coupled to an ultrahigh resolution mass spectrometer (Fusion Lumos tribrid mass spectrometer, Thermo Scientific) to quantify known (derivatized) metabolites. The data files were processed using Skyline (Henderson et al. 2018; Pino et al. 2020). Here, we focus on the results for malic acid, a compound that is not well-captured with PPL-based solid phase extraction resins.

### Statistics and data availability

Statistical analyses and plotting were done using MATLAB version 2019b. Gridding of data prior to plotting was done with a MATLAB implementation of Data Interpolating Variational Analysis (DIVA for MATLAB, Troupin et al. 2012). Targeted metabolomics data for this project are available at MetaboLights (http://www.ebi.ac.uk/metabolights/) as study accession number MTBLS2356. Environmental data is available from the Biological & Chemical Oceanography Data Management Office (BCO-DMO) (http://lod.bco-dmo.org/id/dataset/861266, and http://lod.bco-dmo.org/id/dataset/3782) and from the BATS FTP data site (http://bats.bios.edu/bats-data).

## Results

### Hydrography and environmental data

Over each annual cycle at the study site, the upper ocean exhibited large changes in stratification and depth of the surface mixed layer, with enhanced thermal stratification and shallow mixed layer depths (< 20 m) characterizing the warm, summer months, a progressive deepening in the fall, and maximum mixed layer depths (100-300 m) in late winter / early spring. The onset of warming occurred in April in each year of the project with the mixed layer depth shoaling to < 20 m by June (Figure 1). The depth of the chlorophyll maximum (CM) measured by a fluorometer ranged from 80 to 100 meters (Supplemental Figure 1).

**Figure 1.**
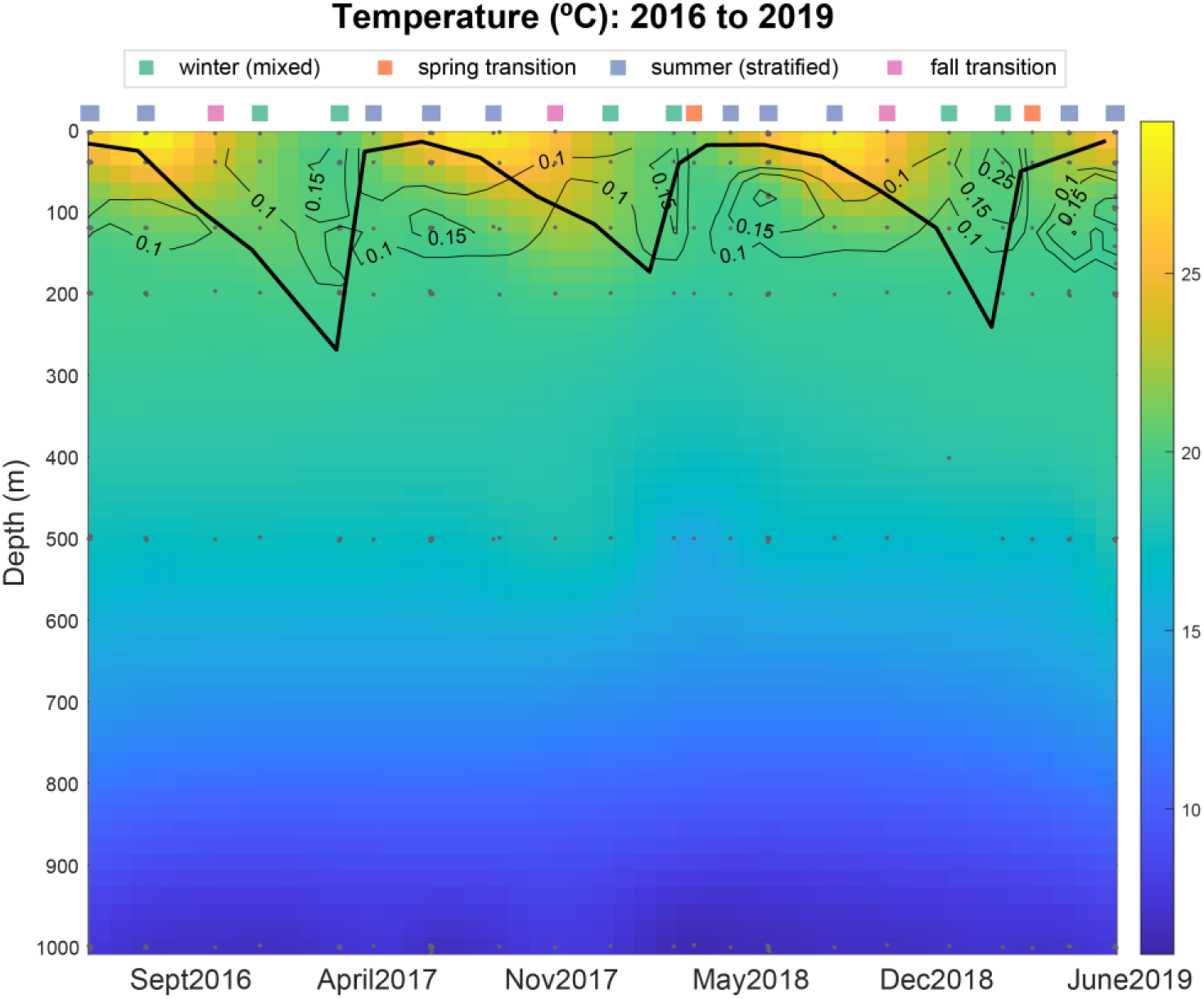
Water temperatures in the upper 1000 meters of the water column from July 2016 through July 2019, the depth and time range of samples that were collected during this project. The grey dots show the locations where discrete samples were collected. The colored boxes at the top indicate the season in which the samples were collected. The contour plot contains fluorescence values. The black line is the mixed layer depth.

We partitioned the upper 1000 m into vertical layers which facilitated statistical comparisons of metabolite concentrations across seasonal and interannual boundaries. From a total of 372 samples, 75% corresponded to the summer (stratified) period when we occupied the station for multiple days during dedicated process cruises. Approximately 15% of the samples were collected during the winter (mixed) period, and the remaining 10% were associated with the relatively brief spring and fall transition periods (Supplemental Table 3). Sample collection was evenly distributed throughout the upper 1000 m of the water column, with 25% of samples collected in the surface mixed layer (VZ_0_) which ranged between 20 to 270 m over the four year period.

Out of the 95 metabolites available in the analytical methods for this project, 41 compounds passed the quality checks and were present in all four years of the dataset. The majority of metabolites not measured have a low extraction efficiency in seawater (Supplemental Table 2) and therefore their absence was not unexpected. Of the metabolites we detected, most exhibited higher concentrations near the surface and decreasing concentrations with increasing depth in the water column (Supplemental Figure 2). Thus, correlations between the concentration of each metabolite and other environmental variables generally revealed significant negative correlations with depth and nutrients, and positive correlations with temperature, bacterial production, and the concentration of bulk organic carbon (measured as TOC or DOC) (Supplemental Figure 3). Metabolites including glyphosate, syringic acid, and cyanocobalamin were present in all four years of the project, were highly variable, and had no significant correlations with measured environmental parameters. In the sections that follow we focus our discussion on metabolites with repeatable patterns throughout the year and repeatable diurnal patterns. We also compare the temporal variability in metabolite concentrations at BATS to the geographical variability of metabolite concentrations sampled along a latitudinal transect between 38°S and 55°N in the western Atlantic Ocean (Johnson et al. accepted, 2022).

### Seasonality of metabolites: mixed versus stratified season

We had sufficient samples from two seasons, mixed and stratified, to allow for rigorous statistical comparisons of metabolite concentrations in the photic zone, the subphotic zone, and the deep ocean (Supplemental Table 3). Differences in metabolite concentrations were most pronounced when comparing the photic zone between the winter (mixed) and summer (stratified) seasons (*p*-values < 0.05, Kruskal-Wallis test) (Table 1, Supplemental Figure 4). The nucleosides adenosine and xanthosine were elevated in the winter, as was S-(5’-adenosyl)-L-homocysteine, the vitamins pantothenic acid (B_5_) and riboflavin (B_2_), and the vitamin precursor desthiobiotin. The metabolic precursor to thiamine (vitamin B_1_), 4-methyl-5-thiazoleethanol (HET), was lower in the winter. While thiamine was measured at BATS during this project, it did not show a seasonal difference. A heterogenous set of compounds, including amino acids and the nucleoside inosine were also higher in the summer.

**Table 1.** Seasonal differences in dissolved metabolites found in the photic zone (VZ_0_, VZ_1_, and VZ_2_, see Supplemental Table 3 for details about the parameters used to define the vertical zones).

| Higher in winter<br>Lower in summer | Higher in summer<br>Lower in winter |
| --- | --- |
| Adenosine | 4-Aminobenzoic acid |
| Desthiobiotin | 4-methyl-5-thiazoleethanol (HET) |
| Pantothenic acid (vitamin B<sub>5</sub>) | D-Ribose 5-phosphate |
| Riboflavin (vitamin B<sub>2</sub>) | Glutathione |
| S-(5'-adenosyl)-L-homocysteine | Inosine |
| Xanthosine | Leucine |
|  | Malic acid |
|  | Taurocholic acid |

### Seasonality of specific metabolites

The set of metabolites with repeatable seasonal patterns over the four years include pantothenic acid, taurocholic acid, and tryptophan. Each showed higher concentrations in the upper 300 meters of the water column.

*Pantothenic acid* - Concentrations ranged from below detection (0.7 pM) to 34.5 pM, with the lowest concentrations at the onset of the stratified period (Figure 2a). In the winter (mixed) period, pantothenic acid accumulated over the upper 300 m, although its concentrations in the winter of 2017/2018 were anomalously low compared to the two other winter periods. Pantothenic acid concentrations integrated within the mixed layer showed a recurring annual pattern of decreased stocks during stratified periods when the mixed layer depth had shoaled to < 20 m (May, July, and September), whereas stocks increased when the mixed layer depth extended deeper than 150 m (January, February, March, and April) (Figure 2b).

**Figure 2.**
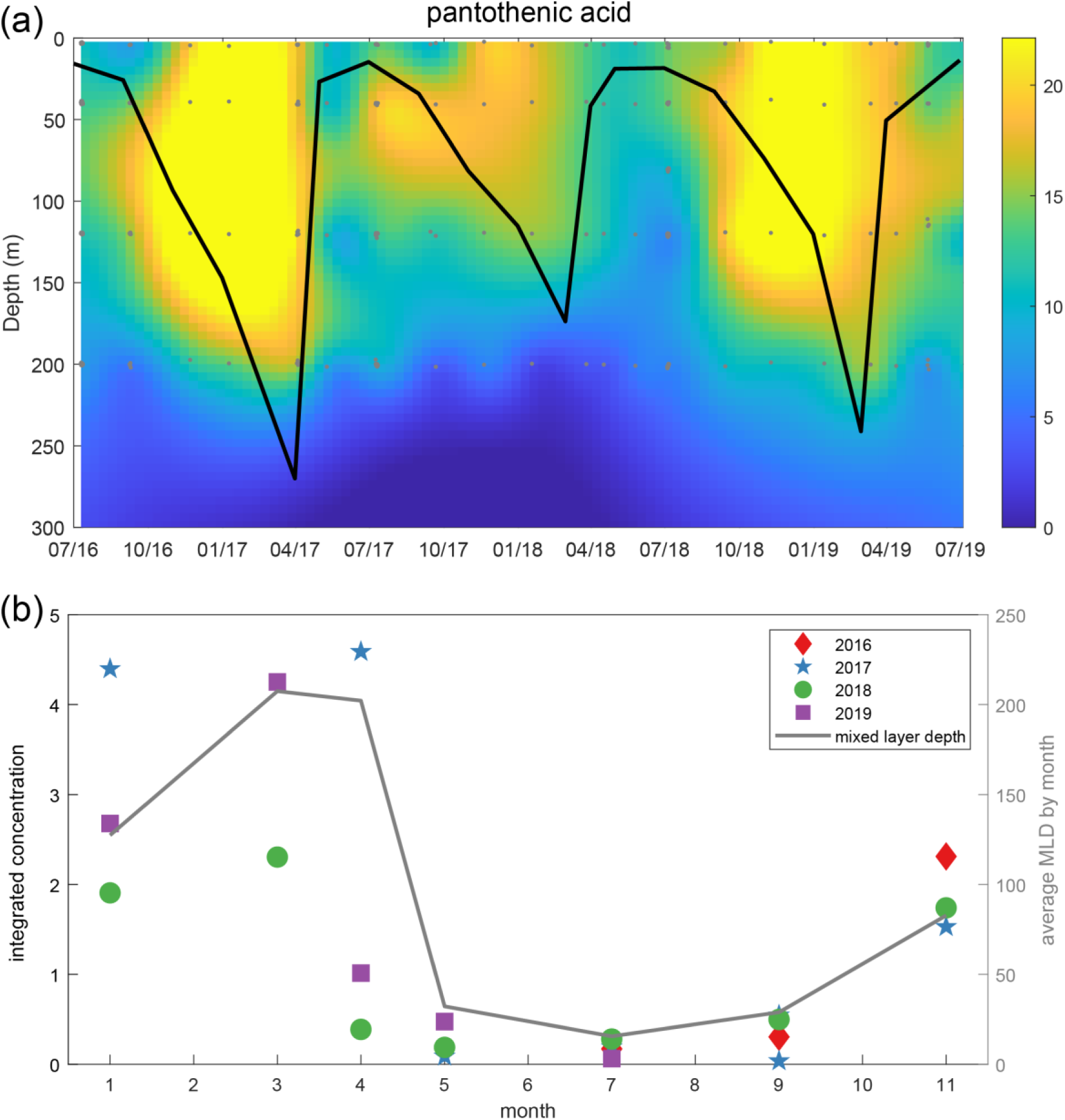
Concentrations of dissolved pantothenic acid in the water column from July 2016 through July 2019, (a) shows pantothenic acid (in pM) in the upper 300 meters of the water column. The grey dots represent discrete samples and the black line is the mixed layer depth over the sampling period, (b) presents the amount of pantothenic acid integrated to the mixed layer depth (in units of µM m-2) and normalized to the integration depth. Values were grouped by month and year before averaging in order to present data from each year over a 12-month annual cycle. The gray line is the averaged mixed layer depth.

*Taurocholic acid* – Mean concentrations of taurocholic acid reached a relative maximum of 2.2 ± 0.8 pM in VZ_0_ during thermal stratification in July (Supplemental Figure 5). Below 100 m, concentrations of taurocholic acid were lower than 1 pM and a weak seasonal pattern with increased concentrations during the summer (stratified) periods in the deep ocean was observed (Supplemental Figure 4).

*Tryptophan*- Regular seasonal variability was also observed for tryptophan with the greatest concentrations in July; however, unlike taurochloric acid the temporal variability was most pronounced between 40 and 120 meters where the mean concentration was 2.6 ± 3.3 pM (Figure 3).

**Figure 3.**
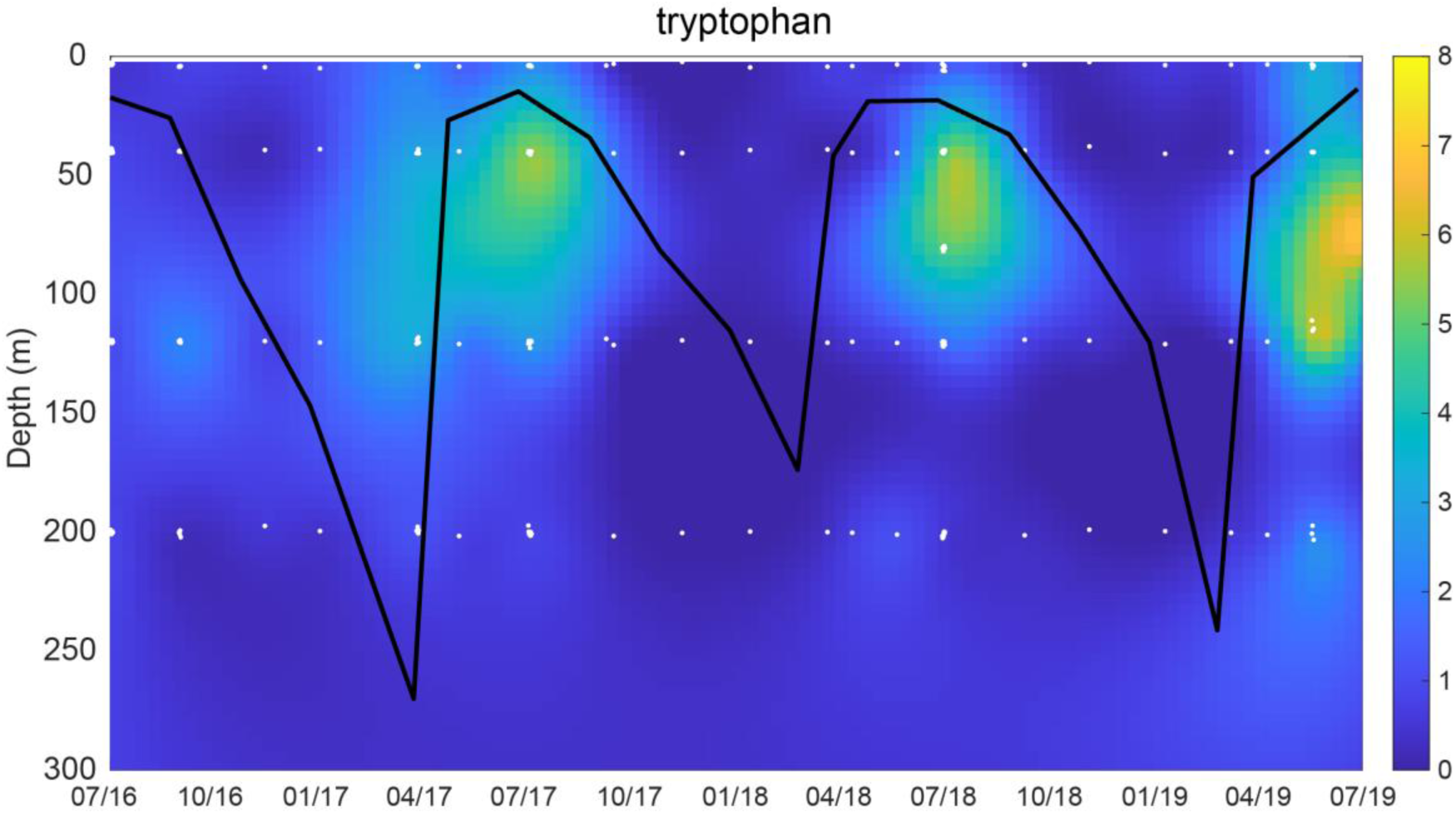
Dissolved tryptophan from July 2016 through July 2019 in the upper 300 m of the water column. The black line is the mixed layer depth. Colors represent the concentration of tryptophan in pM.

### Temporal and vertical variability of metabolites during summer stratification

Metabolite samples were collected every six hours over three days during the July process cruises in each year of the study.

*Riboflavin* - Concentrations of riboflavin demonstrated consistent diel variability in each year of the study with concentrations being lowest in shallowest depths during the mid-day when sampling was coincident with the highest PAR values (Figure 4). The difference in riboflavin concentrations in the daytime (based on sunrise and sunset times at BATS) versus nighttime samples was statistically significant (*p*-value < 0.01). The exception was in July 2019 when riboflavin concentrations were maximal deeper in the water column (between 80 and 100 m) with values mostly below detection in the surface samples (Supplemental Figure 6). Thus, riboflavin presented significant differences on both a diel cycle and a seasonal cycle, as noted in the previous section.

**Figure 4.**
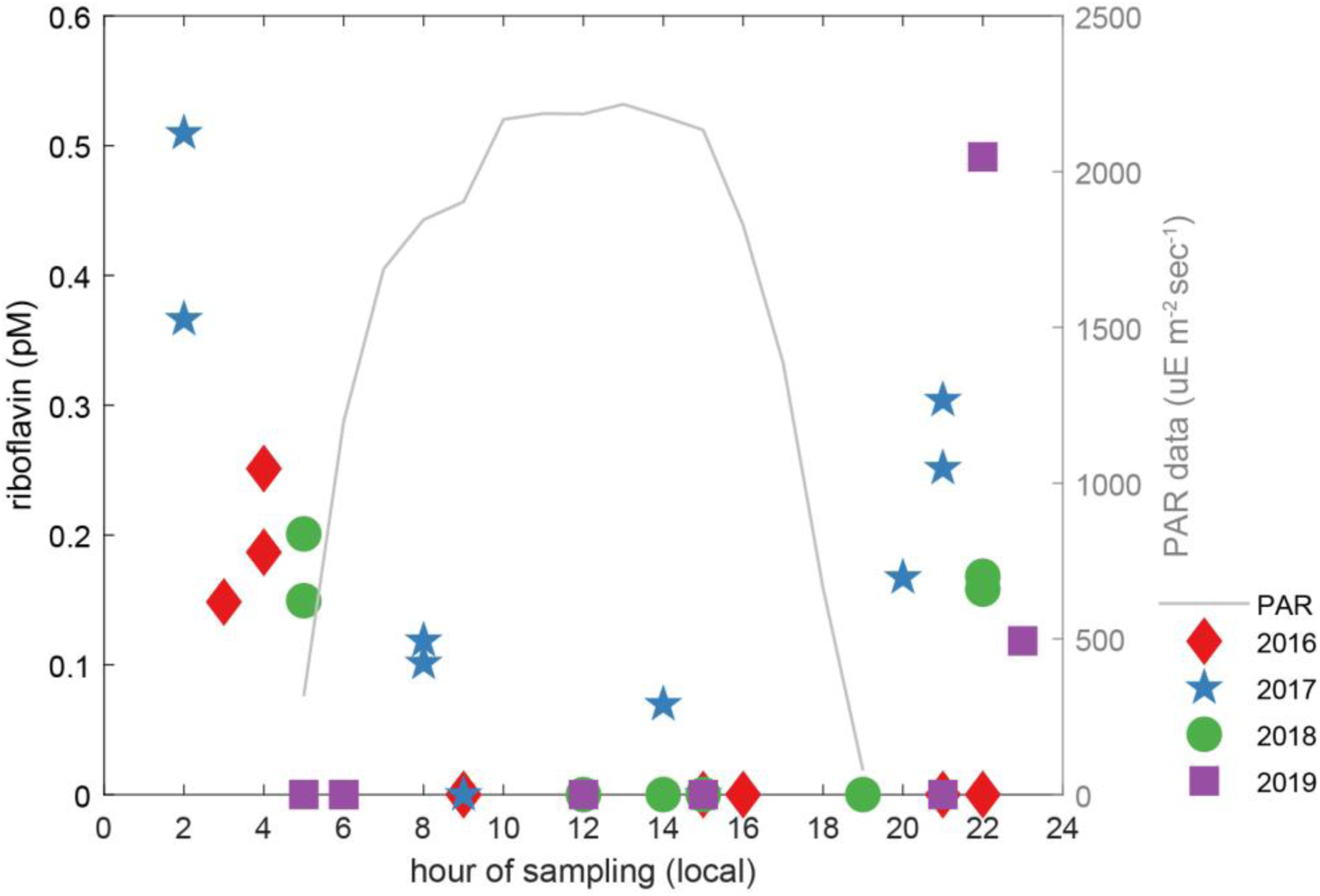
Concentration of dissolved riboflavin concentrations (pM) in the surface water samples from VZ_0_, sampled over 24 hour periods during the July cruises in all years of the project. The PAR data shows the average light levels by hour of the day from the 2017 cruise. The x-axis shows local time.

*Malic acid* – Generally higher concentrations of malic acid (∼500 pM) were observed in the surface ocean and decreased with depth (Figure 5). However, the attenuation patterns over depth were not consistent from year to year. For example, in 2016 the measured concentration for malic acid was greatest at 200 m. One caveat to consider for this metabolite is the extraction efficiency of malic acid with PPL cartridges is low (0.7%, Johnson et al. 2017) and therefore, we did not correct the measured concentrations given the error associated with the correction at low extraction efficiencies. The actual concentrations are likely much higher than those shown in Figure 5a. During the July 2019 cruise we used a pre-extraction chemical derivatization method described in Widner et al. (2021) to validate our observations of malic acid. This method derivatizes the -OH group on malic acid thereby enhancing its extraction with the PPL resin. Using this method we quantified higher concentrations of malic acid approaching 2000 pM at the surface and decreasing with depth to a minima at 600 m. The increased sampling resolution made possible with the benzoyl chloride derivatization method also revealed a deep secondary mesopelagic maximum ranging between 600 – 1700 pM at a depth range of 200 to 600 meters (Figure 5b).

**Figure 5.**
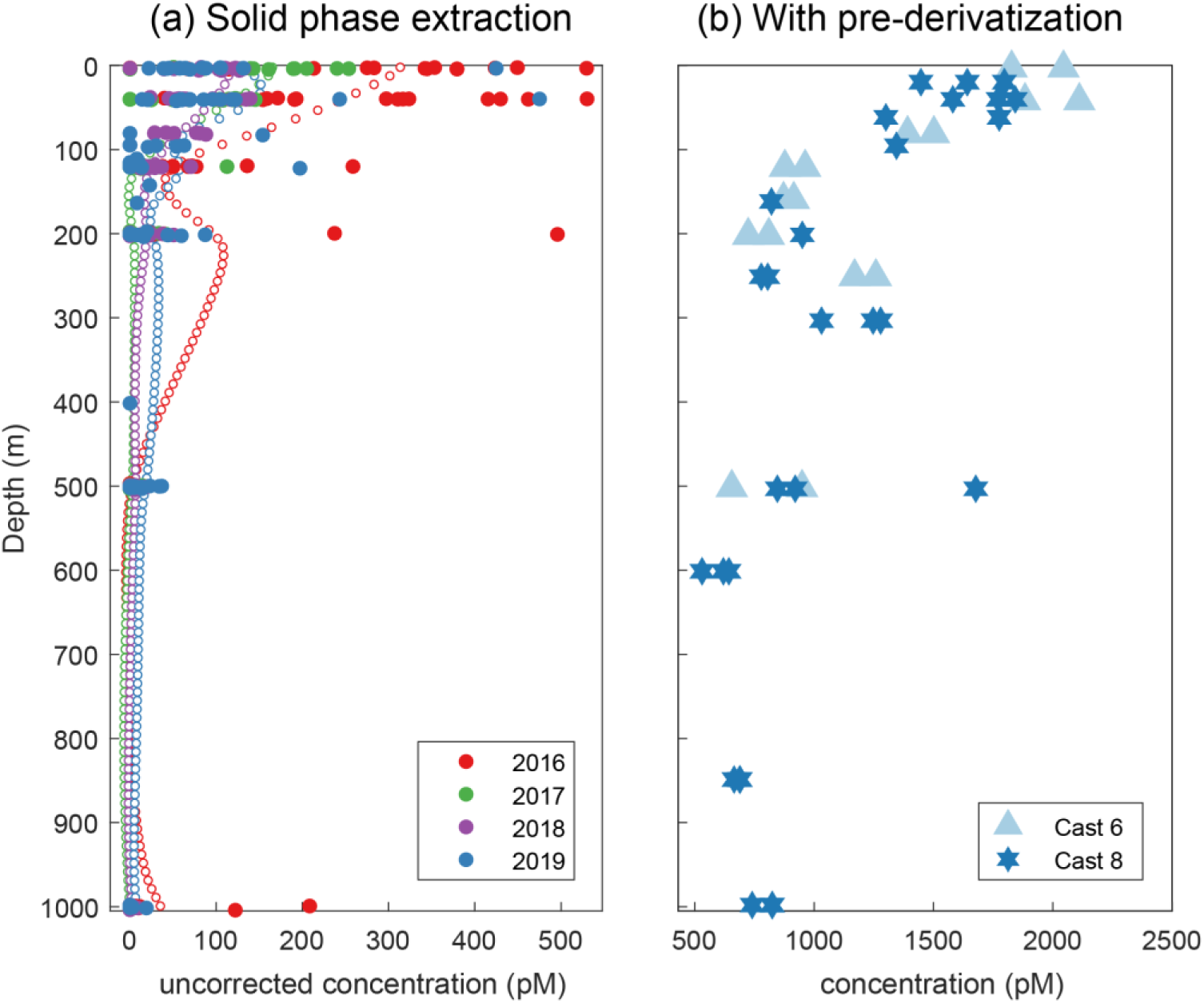
Two methods for sample preparation were used to measure dissolved malic acid during this project. (a) shows data from solid phase extraction using a PPL resin that was used for all four years of samples collected, and (b) shows malic acid captured with a benzoyl chloride derivatization used prior to PPL extraction in the July 2019 cruise. The data in (A) are the discrete samples (filled dots) and interpolated profiles generated using DIVA gridding (open circles). The concentration of malic acid with the solid phase extraction method in (A) has not been corrected for the extraction efficiency of malic acid.

Additional examples of metabolites that showed generally higher concentrations in the surface and decreased throughout the euphotic zone include 4-hydroxybenzoic acid and 5′-methylthioadenosine (MTA). However, there was significant interannual variability in the shape and magnitude of the depth profile for these metabolites (Supplemental Figure 6). 4-hydroxybenzoic acid was higher throughout the upper water column, except in 2018 where it was below detection in all samples collected at the surface. For MTA, the magnitude and location of the highest concentrations varied by year. In 2017, samples from depths less than 10 m had the highest concentrations of MTA, while the samples collected in 2018 and 2019 near the deep chlorophyll maximum presented elevated concentrations (Supplemental Figure 6).

### Comparing temporal versus spatial variability in metabolites

Data from the current study reveal temporal variability in metabolite concentrations over a four-year period in one geographic location. To compare this temporal variability to geographic variability, we calculated the relative standard deviation (RSD, standard deviation divided by the mean) for eleven metabolites found both in the vicinity of BATS and in a study conducted in the western Atlantic Ocean between 38°S and 55°N latitude in which samples were collected in the austral fall and boreal late summer to early fall of 2013 (Johnson et al. accepted, 2022). For the latitudinal study, we restricted the analysis to samples collected from the upper 1000 meters of the water column to allow an explicit comparison between the two datasets. The metabolites in this comparison are all observed in picomolar concentrations (Supplemental Figure 2), and by calculating the RSD we can compare variability in space and time without considering differences in mean values. When RSD values from the BATS site are plotted against those from the latitudinal transect, most of the metabolite abundances showed patterns of higher variability in space compared to the variability in time (Figure 6; symbols below the 1:1 line). Only the amino acid phenylalanine showed higher temporal variability compared to the latitudinal transect, and three metabolites (4-aminobenzoic acid, caffeine, and tryptophan) had the same degree of variability in time and space. The remaining metabolites showed greater degree of variability across geographic location. As a comparison, the RSD of dissolved organic carbon at the BATS site was 15.6% compared to 22.3% in the samples collected along the latitudinal transect in the western Atlantic Ocean.

**Figure 6.**
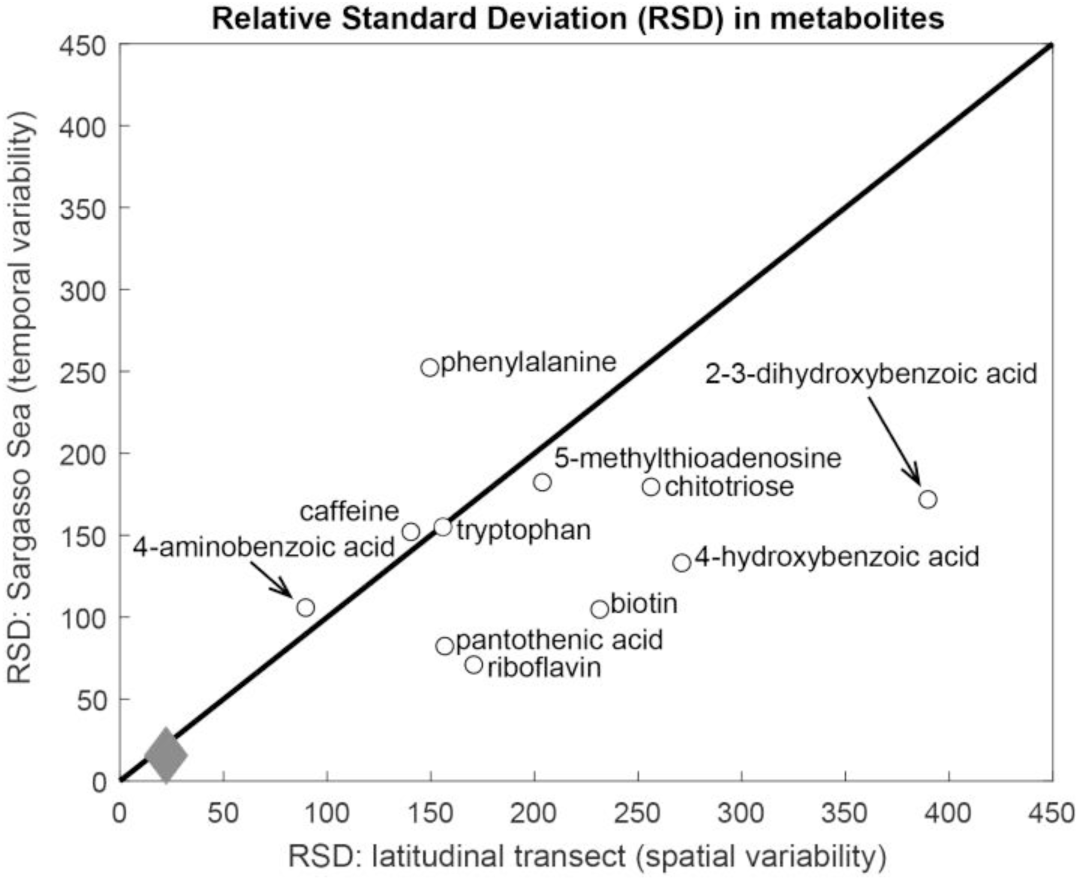
Relative standard deviation (RSD) in dissolved metabolite concentrations from 2016 to 2019 in the Sargasso Sea (y-axis) compared with data from a latitudinal transect in the western Atlantic Ocean (x-axis, Johnson et al. accepted, 2022). The black line is the one-to-one line where the variability in time (Sargasso Sea) matches the variability in space (western Atlantic Ocean transect). The arrows are used to connect metabolite names with data points when crowding would prevent names from appearing adjacent to its datapoint. The gray diamond is the RSD of bulk concentrations of total/dissolved organic matter in each dataset.

## Discussion

The inventory of marine DOC represents the largest reservoir of exchangeable reduced carbon in the ocean. The biological and physical dynamics that control the sources and sinks of marine DOC over space and time play a critical role in the global carbon cycle. Our understanding of the changes in bulk DOC has evolved over the years (Carlson and Hansell 2015), and here we provide details on a subset of small organic metabolites that comprises a portion of the myriad of compounds within dissolved organic matter. Our four years of data provides evidence of recurring temporal patterns of compounds that systematically vary on time scales of an annual cycle (month to month variability), on time scales of seasons (winter versus summer), and a compound that varies over a diel cycle in summer.

### An annually repeatable pattern: pantothenic acid

The water-soluble vitamin pantothenic acid (vitamin B_5_) is the clearest example of a compound with an annual pattern that repeats regularly in each of the four years at BATS. Pantothenic acid was present in three-quarters of the samples collected in the upper 300 m, therefore reflecting the origin of its name from the Greek ‘pantos’ meaning from everywhere. Its concentration, however, was not constant throughout the year. We measured the lowest concentrations in the late spring/early summer periods when thermal stratification was greatest and highest concentrations during periods of vigorous mixing in winter and early spring as the mixed layer depth increased. Pantothenic acid was discovered in the 1930s as a growth factor for yeast (Williams et al. 1933), and subsequent work has shown that pantothenic acid is incorporated intracellularly into CoA, a cofactor used in common metabolic pathways including lipid synthesis, processing of fatty acids, and the tricarboxylic acid cycle (Leonardi et al. 2007; Novelli 1953). Microbial production of pantothenic acid can exceed demand for the compound (Jackowski and Rock 1981), which may lead to its release from cells. While microorganisms require pantothenic acid, details on which organisms produce dissolved pantothenic acid and why it is released extracellularly in marine systems are sparse. In laboratory cultures, three strains of the abundant marine phytoplankton *Prochlorococcus* have been shown to release pantothenic acid to the surrounding media (Kujawinski et al. submitted, December 2022); at BATS the distribution of *Prochlorococcus* is seasonally variable (Olson et al. 1990) with maximum concentrations usually between 60 and 80 m (DuRand et al. 2001). Furthermore, in heterotrophic marine organisms, genetic information reveals that the abundant SAR86 group lacks a putative metabolic pathway to produce pantothenic acid (Dupont et al. 2012). While broad statements about sources of pantothenic acid in marine environments are not yet possible, Liu et al. (2022) proposed that some groups of bacterioplankton may shift their metabolisms as the mixed layer depth shoals following deep convective mixing resulting in enhanced scavenging of pantothenic acid in the surface 200 m at BATS. Clearly, additional research is needed to define the balance of sources and sinks that lead to the repeatable pattern of lower pantothenic acid concentrations during the summer period in the Sargasso Sea.

### Seasonal shifts in the balance of dissolved metabolites

Our *in situ* dissolved metabolite concentration data reveal changes in the standing stock of a metabolite, or the balance between changes in metabolite production and consumption. An increase in the amount of a metabolite measured in the water column can indicate an increase in its production or a decrease in its consumption, or a change in the physical transport of metabolites in and out of the ecosystem. We cannot separate production and consumption without additional information. With these caveats in mind, we next consider the metabolites that had relatively higher concentrations in the summer (stratified) period compared to the winter (mixed) period.

The seasonal pattern of bulk DOC dynamics and its role as an export pathway in the biological carbon pump has been well established at BATS (Carlson et al., 1994, Hansell and Carlson 2001). Briefly, the DOC concentrations increase in the euphotic zone as the water column stratifies in late spring or early summer and remain elevated until the mixed layer extends deeper than the euphotic zone in the winter or early spring. During winter convective mixing to depths between 200 and 300 m, DOC is homogeneously redistributed throughout the mixed layer and as a result a portion of the seasonally accumulated DOC is exported from the euphotic zone into the mesopelagic zone (Carlson et al. 1994; Hansell and Carlson 2001). DOC accumulation within the euphotic zone results from a relative imbalance between DOM production processes and heterotrophic bacterial consumption (Carlson et al. 1996). Factors that can affect bacterial production and control the accumulation of DOM include potential inorganic limitation of heterotrophic bacterioplankton production (Cotner et al. 1997; Thingstad et al. 1997), the production of recalcitrant organic compounds (Aluwihare and Repeta 1999) that resist rapid microbial remineralization, and the composition and varying metabolic potential of the resident microbial communities within the euphotic zone (Carlson et al. 2004). At BATS this imbalance results in the accumulation of dissolved combined neutral sugars (Goldberg et al. 2009) and total combined amino acids (Liu et al. 2022) during the summer stratified period. In the winter, seasonal mixing redistributes DOC that accumulated in the surface to deeper in the water column; organic carbon at deeper depths is then converted to inorganic carbon via microbial respiration as the water column re-stratifies (Hansell and Carlson 2001).

The present study showed significant differences in dissolved metabolites that accumulate within the mixed layer during the winter (mixed) period compared to the summer (stratified) period. While some metabolites revealed seasonal patterns, as was expected based on the known variability in bulk organic carbon, other metabolites did not reveal regular patterns, which emphasizes the need to quantify changes in individual compounds. For some compounds, these differences were also evident at deeper depths (500 - 1000 m, Supplemental Figure 2), though the number of seasonally-varying compounds was highest at shallower depths (Supplemental Figure 4). The compounds with higher concentrations in the winter were predominantly vitamins, in contrast to amino acids, nucleic acid precursors, and other metabolites that were most pronounced in the summer.

The excess S-(5’-adenosyl)-L-homocysteine (SAH) in the water column during the winter period could be an indication of greater release by cells seeking to avoid the negative effects of higher intracellular levels of SAH. SAH is produced when S-adenosyl-L-methionine (SAM) donates methyl groups to molecules such as DNA, RNA, or proteins (Parveen and Cornell 2011); however, the cellular accumulation of SAH inhibits this methylation. Halophilic cyanobacteria can generate SAH during the synthesis of compatible solutes used to endure increased salt concentrations (Sibley and Yopp 1987), and SAH can be converted to adenosine and homocysteine by prokaryotic cells (Shimizu et al. 1984). The higher levels of adenosine also observed during the winter at BATS could represent an additional sink of SAH in the photic zone during the winter. In *Prochlorococcus*, the balance between SAH and SAM inside the cells showed statistically significant differences across strains and growth conditions, indicating that broad statements about the production of SAH in marine phytoplankton are not possible (Kujawinski et al. submitted, December 2022). In SAR11, an abundant cosmopolitan marine heterotrophic alphaproteobacteria, the transcription of genes producing SAH are enhanced under sulfur-limited growth (Smith et al. 2016). This is of particular relevance given SAR11’s inability to use oxidized forms of sulfur such as sulfate and its growth requirement for exogenous reduced sulfur (Tripp et al. 2008). At BATS, the lowest levels of the organic sulfur compounds dimethylsulfide (DMS) and dimethylsulfoniopropionate (DMSP) occur during the winter (Levine et al. 2016). Hence, if SAR11 has an inadequate supply of reduced sulfur in the winter at BATS, it could increase its transcription of genes producing SAH and thereby release increased amounts of SAH into the water column as we observed in our data.

The biochemical reactions described above whereby SAH is produced during the methylation of molecules, also produce MTA (Parveen and Cornell 2011). We have measured MTA in the dissolved phase in both the north Atlantic (Johnson et al. accepted, 2022), and in laboratory cultures with *Ruegeria pomeroyi* (Johnson et al. 2016) and *Prochlorococcus* MIT9313 (Kujawinski et al. submitted, December 2022). Intracellularly, MTA is significantly less abundant in *Thalassiosira pseudonana* when cobalamin concentrations are low (Heal et al. 2019), which indicates a link between internal MTA levels and conditions experienced by phytoplankton. At BATS, dissolved MTA concentrations peak at different depths in the water column, in different years, a pattern that was not correlated to any single environmental factor. Furthermore, the MTA levels did not show a significant seasonal differences indicating further investigation is required to determine the environmental conditions that control the distribution of MTA in the water column.

Two vitamins (pantothenic acid and riboflavin) and one vitamin precursor (desthiobiotin) showed higher concentrations in the mixed layer during the winter mixed period. The production and consumption of vitamins occurs in both phytoplankton and heterotrophic bacteria (Koch et al. 2012; Rodionov et al. 2003; Sañudo-Wilhelmy et al. 2014; Warren et al. 2002), and vitamins are exchanged within microbial communities (Joglar et al. 2021; Wienhausen et al. 2022a; Zoccarato et al. 2022). Vitamin precursors may enable cellular growth in the absence of the vitamin itself, as has been observed in some, but not all, bacterial cells tested for the ability to use desthiobiotin in the absence of biotin (Wienhausen et al. 2022b). In addition, many species of eukaryotic phytoplankton require external sources of vitamins (Croft et al. 2005; Croft et al. 2006), and eukaryotic picophytoplankton are prevalent in the winter period at BATS (Giovannoni and Vergin 2012) and presumably consume vitamins. However, the winter mixed season is the time when we see relatively higher vitamin concentrations which contradicts the pattern that would be expected if consumption of vitamins by eukaryotic phytoplankton were the dominant controlling factor. Furthermore, while light and increased temperature can cause degradation of vitamins (Carlucci et al. 1969; Gold et al. 1966), only riboflavin revealed significantly lower concentrations in the day compared to night, suggesting that the accumulation of the other vitamins such as pantothenic acid are not directly controlled by lower light levels in the winter.

The higher levels of 4-methyl-5-thiazoleethanol (HET), a precursor to thiamine (vitamin B_1_), is the one vitamin-related compound that showed higher levels during the summer. Auxotrophy for thiamine, the condition where cells require an external source for a compound, is genetically widespread in marine microorganisms (Paerl et al. 2018). While thiamine did not show seasonal differences in the photic zone, our data show higher levels of HET during the summer. Although HET is a precursor for thiamin, bacteria such as *Ostreococcus* (Paerl et al. 2015) and *Emiliana huxleyi* (McRose et al. 2014) that are given HET in lieu of thiamine are not able to grow. There are complex and competing interactions between autotrophic and heterotrophic microorganisms that control the dynamics of vitamins in seawater; thus, additional knowledge is required before we can confirm which factors drive the seasonal balance between production and consumption of vitamins.

The remaining compounds with elevated levels during the summer are select amino acids, nucleic acid precursors, and several miscellaneous compounds. Given the diversity of these compounds, constructing a single hypothesis to explain their distribution over the annual cycle is challenging. Leucine and isoleucine are among the amino acids that are energetically costly to produce (McClelland and Montoya 2002; Yamaguchi et al. 2017) which may indicate conditions in the summer enable relatively higher production. Furthermore, extracellular enzyme activity, especially peptidase activity that cleaves proteins into free amino acids, has been shown to increase with warmer summer temperatures leading to excessive amino acid production relative to consumption (Piontek et al. 2014). Inosine and 4-aminobenzoic acid are among the known compounds excreted by copepods (Maas et al. 2020), which suggests that seasonal changes in the community composition (Blanco-Bercial 2020) or physiology of the vertical migrating zooplankton at BATS could be a source of these metabolites in the mixed layer. D-ribose 5-phosphate, malic acid, and taurocholic acid are also relatively higher in the summer compared to the winter, although the reasons remain unclear.

### Diel shifts in riboflavin (vitamin B_2_) during the summer

The subtropical north Atlantic Ocean experiences higher incoming solar radiation than the temperate and polar latitudes. Riboflavin, which showed significant decreases in concentration during the daylight hours in the surface water masses, was the only measured organic compound responding to daily changes in light. Riboflavin is a water-soluble vitamin that is a precursor for flavin mononucleotide or flavin adenine dinucleotide, cofactors involved in electron transfer, and has been widely-studied since it was first isolated in the 1880s (Eggersdorfer et al. 2012). Riboflavin is subject to direct photodegradation due to its highly-conjugated, aromatic structure, and the history of research on the sensitivity of riboflavin to light dates to the 1930s when the first oxidation products of riboflavin were isolated (Warburg and Christian 1932). The loss of riboflavin occurs at wavelengths of light between 350-520 nm, with the highest levels of damage in the narrower window between 415-455 nm (Ahmad et al. 2006). The interactions between light and riboflavin also vary as a function of pH (Ahmad et al. 2004), ionic strength (Ahmad et al. 2016), and temperature (Sattar et al. 1977). In addition to its role in metabolism, riboflavin can also act as a quorum-sensing molecule (Rajamani et al. 2008). Our dissolved riboflavin concentrations, from below detection to 0.6 pM, are at the low end of the dynamic range of existing data, although coastal seawater from the North Sea had even lower concentrations at less than 0.04 pM (Bruns et al. 2022). Along a north-south transect in the western Atlantic Ocean, Johnson et al. (accepted, 2022) measured values averaging 0.3 pM (range 0 to 12 pM) with the highest concentrations found between 50 and 150 meters, with the exception of the northernmost station where values approached 12 pM at 1 m. These values are comparable to dissolved riboflavin concentrations off the coast of California (Sañudo-Wilhelmy et al. 2012). In contrast, in a coastal inlet in the northwest coast of the United States, Heal et al. (2014) found riboflavin had the highest concentrations of the B vitamins, with values ranging from 40 to 120 pM. In laboratory cultures, riboflavin is released by *Synechococcus* (Fiore et al. 2015) and *Thalassiosira* (Kujawinski et al. 2017; Longnecker and Kujawinski 2017), but is not released by *Prochlorococcus* (Kujawinski et al. submitted, December 2022). Coral reefs (Weber et al. 2022) and sponges (Fiore et al. 2017) are also sources of riboflavin to the marine environment.

The daytime decrease in riboflavin concentrations could be due to the response of riboflavin to light, consumption by the *in situ* microbial community, or a combination of both processes. In the Pacific Ocean, intracellular concentrations of riboflavin reached maximal levels at the end of the day (Boysen et al. 2021). Furthermore, while riboflavin is known to react to light, there are certainly other, unknown, compounds in the euphotic zone that exhibit this behavior. To our knowledge, there are no organisms known to be auxotrophic for riboflavin nor has a daily cycle in riboflavin production been examined. The daily decrease in riboflavin in the surface ocean has implications for the microorganisms who rely on production of vitamins by other organisms because excess dissolved riboflavin is only present at night.

### Additional compounds of interest

Dissolved malic acid concentrations were higher in the summer in the upper euphotic zone, in VZ_0_. Malic acid is an intermediate in the citric acid cycle and is produced in the first step in the Hatch-Slack, or C_4_ carbon fixation, pathway. The C_4_ carbon fixation pathway is less common and on land is dominated by grasses (Sage 2016). In marine ecosystems there is equivocal evidence for a C_4_ photosynthetic pathway in diatoms (Mackey et al. 2015), and diatoms with the C_4_ photosynthetic pathway may use it to dissipate excess light energy rather than to fix carbon dioxide (Haimovich-Dayan et al. 2013). The Tara Oceans dataset revealed low transcript levels for the enzymes of the C_4_ pathway (Pierella Karlusich et al. 2021), supporting the possibility that the higher levels of malic acid in the surface ocean are due to exudation by eukaryotic phytoplankton. Malic acid is one of multiple organic compounds measured within marine aerosols collected in the western Arctic Ocean (Kawamura et al. 2012) and western Pacific Ocean (Kawamura and Sakaguchi 1999). Thus, we cannot rule out the possibility of an atmospheric source of malic acid to the surface ocean.

The pattern of 4-hydroxybenzoic acid in the water column is an example of a metabolite with irregular vertical and temporal variability during our multi-year project. 4-hydroxybenzoic acid was completely absent from the shallowest samples in 2018, yet was present deeper in the water column at similar concentrations for all years. Furthermore, the return of 4-hydroxybenzoic acid to the surface waters in 2019 indicates a transient decoupling between the production and removal processes in the upper water column. In laboratory cultures, cyanobacteria release 4-hydroxybenzoic acid (Fiore et al. 2015; Kujawinski et al. submitted, December 2022) or use it as a carbon source (Mou et al. 2007), which complicates defining sources and sinks within the water column. In shallow reef habitats, marine sponges remove 4-hydroxybenzoic acid from the water column, likely due to the actions of the microbial community residing within the sponges (Fiore et al. 2017). The presence of 4-hydroxybenzoic acid in the water column will have unknown effects on the microbial community as it can both stimulate and inhibit prokaryotic activity in a manner that varies for different microbial species (Czerpak et al. 2001; Kamaya et al. 2006). Our previous work (Liu et al. 2020b) demonstrates that we can analytically separate the isomers 3-hydroxybenzoic acid (*m*-hydroxybenzoic acid) and 2-hydroxybenzoic acid (*o*-hydroxybenzoic acid), from 4-hydroxybenzoic acid (*p*-hydroxybenzoic acid); however, these isomers were not analyzed during this project. This is relevant because the effects of 2- and 3-hydroxybenzoic acid differ from 4-hydroxybenzoic acid on the microbial community, as 2-hydroxybenzoic acid strongly simulates growth, 3-hydroxybenzoic acid inhibits growth, and 4-hydroxybenzoic acid weakly stimulates growth (Czerpak et al. 2001). Thus, the biological community responds differently to each isomer, underscoring the care that must be taken with respect to measuring structural isomers in marine metabolomics.

### Scale of variability in metabolites

By comparing data from this multi-year sampling with our previous data from the western Atlantic Ocean (Johnson et al. accepted, 2022), we find that the variability in dissolved metabolite abundance across space is generally greater than the variability over time. The greater spatial variability in metabolites is consistent with the expected variability in biological communities over large geographic distances. Furthermore, the variability in the concentrations of individual dissolved metabolites is considerably higher than the variability in bulk organic carbon concentrations, emphasizing that changes in bulk dissolved organic matter can mask changes in the components that make up the pool of dissolved organic matter. The relatively higher variability in metabolites along a latitudinal transect indicates that a single time-series station cannot serve as a model for the global ocean. However, the repeated sampling at BATS enables us to investigate long-term patterns in metabolites as the ocean encounters future changes in climate. Since the 1980s, the BATS site has become warmer and more acidic (Bates and Johnson 2020). While we cannot yet observe long-term changes in our dataset, in future work we can track metabolites over time to view how specific organic compounds change within the context of a changing climate and the subsequent impact on the marine microbial food web.

## Conclusions

The dissolved metabolites measured in the northwestern Sargasso Sea during this multi-year study are central carbon metabolites and examining their variability in time provides insight into the processes that underlie chemical variability in a marine ecosystem. These time series data reveal recurring temporal patterns of known dissolved organic compounds on both diel and seasonal timescales. How the temporal variability of dissolved metabolites is linked the sources and sinks of other biological and biogeochemical variables is the next challenge to determine how marine metabolites will respond to future changes in the marine environment.

## Acknowledgements

We thank Rod Johnson and the team of BATS technicians (especially Julia Matheson, Claire Medley, Zac Anderson, Dominic Smith and Paul Lethaby); the BIOS-SCOPE project team for assistance with sample collection and useful discussions about the project’s results; Katelyn McLeod, Miguel Desmarais, Craig McLean, Keri Opalk for assistance with sample processing; Elisa Halewood for assistance with metadata curation; and Erin McParland and Noah Germolus for providing comments on the manuscript. The efforts of the captains, crew, and marine technicians of the R/V *Atlantic Explorer* are a key aspect of the success of this project. The samples analyzed in this project were exported from Bermuda to the United States under Bermuda Institute of Ocean Sciences (BIOS) export permit numbers SP160704, SP170701, and SP180501. This work was supported by funding from the Simons Foundation International’s BIOS-SCOPE program.

## Supplemental Information

**Supplemental Table 1.** Samples were collected over the period spanning July 2016 to July 2019. Samples were collected from three sites: Hydrostation S (32°10’N, 64°30’W), Bermuda Atlantic Time-series Study station (BATS, 31°40’N, 64°10’W), and east of BATS (AE1916, 32°10’N, 64°13’W) and select cruises had increased sampling frequency when the station was occupied for multiple days. Samples were collected from the surface ocean down to 1000 m (Supplemental Table 3).

| Year | Month | # samples | Station | Cruise |
| --- | --- | --- | --- | --- |
| 2016 | 7 | 56 | Hydrostation S | AE1614 |
| 2016 | 9 | 24 | Hydrostation S | AE1620 |
| 2016 | 11 | 6 | BATS | AE1627 |
| 2017 | 1 | 6 | BATS | AE1701 |
| 2017 | 4 | 27 | BATS | AE1703 |
| 2017 | 5 | 6 | BATS | AE1707 |
| 2017 | 7 | 60 | BATS | AE1712 |
| 2017 | 9 | 3 | BATS | AE1718 |
| 2017 | 9 | 6 | Hydrostation S | AE1719 |
| 2017 | 11 | 6 | BATS | AE1726 |
| 2018 | 1 | 6 | BATS | AE1802 |
| 2018 | 3 | 6 | BATS | AE1809 |
| 2018 | 4 | 6 | BATS | AE1810 |
| 2018 | 5 | 6 | BATS | AE1814 |
| 2018 | 7 | 53 | BATS | AE1819 |
| 2018 | 9 | 6 | BATS | AE1825 |
| 2018 | 11 | 6 | BATS | AE1831 |
| 2019 | 1 | 6 | BATS | AE1902 |
| 2019 | 3 | 6 | BATS | EN631 |
| 2019 | 4 | 6 | BATS | AE1905 |
| 2019 | 5 | 18 | BATS | AE1910 |
| 2019 | 7 | 47 | East of BATS | AE1916 |

**Supplemental Table 2.** Complete set of metabolites within the targeted metabolomics method used in the current project. The extraction efficiency information is from Johnson et al. (2017), supplemented with unpublished data on extraction efficiency for new metabolites added to the analytical method (Swarr et al. unpublished). The concentration data for each metabolite is available at MetaboLights (http://www.ebi.ac.uk/metabolights/) as study accession number MTBLS2356. Metabolites found at in this project are marked with ‘Yes’.

| metabolite | extraction efficiency (%) | Sargasso Sea |
| --- | --- | --- |
| 2,3-dihydroxybenzoic acid | 100.9 | Yes |
| 2,3-dihydroxypropane-1-sulfonate | 0.6 | Yes |
| 3-mercaptopropionic acid | 88.6 | Yes |
| 3-methyl-2-oxobutanoic acid | 10.1 | Yes |
| 3-methyl-2-oxopentanoic acid | 50.1 |  |
| 4-methyl-5-thiazoleethanol | 11.2 | Yes |
| 4-aminobenzoic acid | 18.5 | Yes |
| 4-hydroxybenzoic acid | 88 | Yes |
| 4-methyl-2-oxopentanoic acid | 43.4 |  |
| 4-amino-2-methyl-5-pyrimidinemethanol | 0 |  |
| 4-amino-5-aminomethyl-2-methylpyrimidine | 0 |  |
| 5'-methylthioadenosine | 80.9 | Yes |
| γ-aminobutyric acid | 0 | |
| acetyltaurine | 0 |  |
| adenine | 0 |  |
| adenosine | 6.5 | Yes |
| adenosine 5'-monophosphate | 0.2 |  |
| alpha-ketoglutaric acid | 0 |  |
| arginine | 0 |  |
| aspartic acid | 0 |  |
| glycine betaine | 0 |  |
| biotin | 52.8 | Yes |
| caffeine | 23.5 | Yes |
| chitobiose | 0.3 |  |
| chitotriose | 5.6 | Yes |
| choline | 0 |  |
| ciliatine | 0 |  |
| citric acid | 0.9 |  |
| citrulline | 0 |  |
| cyanocobalamin | 79.1 | Yes |
| cysteine | 0 |  |
| cytosine | 0 |  |
| desthiobiotin | 6.5 | Yes |
| D-glucosamine 6-phosphate | 0 |  |
| D-ribose 5-phosphate | 0.1 | Yes |
| dihydroxyacetone phosphate | 0 |  |
| dimethylsulfoniopropionate | 0 |  |
| ectoine | 0 |  |
| folic acid | 41.2 | Yes |
| fosfomycin | 0 |  |
| fumaric acid | 0 |  |
| glucose 6-phosphate | 0 |  |
| glutamic acid | 0 |  |
| glutamine | 0 |  |
| glutathione | 1.1 | Yes |
| glutathione oxidized | 1.5 | Yes |
| glyphosate | 15.1 |  |
| guanine | 0 |  |
| guanosine | 7.7 | Yes |
| hemin | 0 |  |
| indole 3-acetic acid | 16.8 | Yes |
| inosine | 8.1 | Yes |
| inosine 5'-monophosphate | 0 |  |
| isoleucine | 1.41 | Yes |
| kynurenine | 41.7 | Yes |
| leucine | 3.3 | Yes |
| malic acid | 0.7 | Yes |
| methionine | 0 |  |
| muramic acid | 0 |  |
| n-acetyl glucosamine | 0 |  |
| n-acetyl glutamic acid | 1.1 | Yes |
| n-acetyl muramic acid | 2.8 | Yes |
| nicotinamide adenine dinucleotide | 20.9 |  |
| ornithine | 0 |  |
| orotic acid | 0 |  |
| pantothenic acid | 51.9 | Yes |
| phenylalanine | 39.7 | Yes |
| phosphoenolpyruvate | 0 |  |
| phosphoglyceric acid | 0 | Yes |
| phycocyanobilin | 0 |  |
| proline | 0 |  |
| putrescine | 0 |  |
| pyridoxine | 6.8 | Yes |
| riboflavin | 87.6 | Yes |
| S-(5'-adenosyl)-L-homocysteine | 43.3 | Yes |
| S-adenosyl-L-methionine | 0 |  |
| sarcosine / alanine (isomers) | 0 |  |
| serine | 0 |  |
| sn-glycerol 3-phosphate | 0 |  |
| spermidine | 0 |  |
| succinic acid | 0 |  |
| syringic acid | 32.9 | Yes |
| taurine | 0 |  |
| taurocholic acid | 92.8 | Yes |
| thiamine | 2 | Yes |
| thiamine monophosphate | 0 |  |
| threonine / homoserine (isomers) | 0 |  |
| thymidine | 52.6 | Yes |
| tryptamine | 21.3 | Yes |
| tryptophan | 46.7 | Yes |
| tyrosine | 2.1 | Yes |
| uracil | 0 |  |
| uridine 5'-monophosphate | 0 |  |
| valine | 0 |  |
| xanthine | 0.3 |  |
| xanthosine | 10.4 | Yes |

**Supplemental Table 3.** A total of 372 samples were collected in the time period spanning July 2016 to July 2019. The samples were divided into the vertical zones and four seasons based on the definitions in the table. The winter vs. summer column indicates sufficient samples were collected during both the winter (mixed) period and the summer (stratified) period to enable a seasonal comparison of metabolites. In some vertical zones samples were only collected in one season (e.g., vertical zone 1 only has samples from the summer). No statistical comparisons during the transition periods were conducted due to the low number of samples. In the descriptions ‘sigma’ refers to the sigma-theta density layer, and the following abbreviations are used: MLD, mixed layer depth; CM, chlorophyll maximum; WMW, winter mode layer water.

|  |  |  | # of samples | Separation of samples by season |  |  |  | winter vs. summer |
| --- | --- | --- | --- | --- | --- | --- | --- | --- |
|  |  |  |  | winter, mixed | spring, transition | summer, stratified | fall, transition |  |
| Vertical zone | Description | Layer |  | Season 1 | Season 2 | Season 3 | Season 4 |  |
| 0 | Surface mixed layer | photic layer | 91 | 33 | 4 | 48 | 6 | Yes |
| 1 | MLD to top of CM layer |  | 46 | 0 | 0 | 46 | 0 |  |
| 2 | CM layer |  | 54 | 0 | 1 | 51 | 2 |  |
| 3 | WMW layer | subphotic layer | 64 | 5 | 2 | 53 | 4 | Yes |
| 4 | Base of WMW to sigma = 26.5 | deep | 14 | 3 | 1 | 9 | 1 | Yes |
| 5 | Sigma 26.5 to 26.7 |  | 47 | 6 | 1 | 38 | 2 |  |
| 6 | Sigma 26.7 to 276.9 |  | 2 | 1 | 1 | 0 | 0 |  |
| 9 | Sigma 27.3 to 27.5 |  | 15 | 2 | 0 | 12 | 1 |  |
| 10 | Sigma > 27.5 |  | 39 | 7 | 2 | 28 | 2 |  |

**Supplemental Figure 1.**
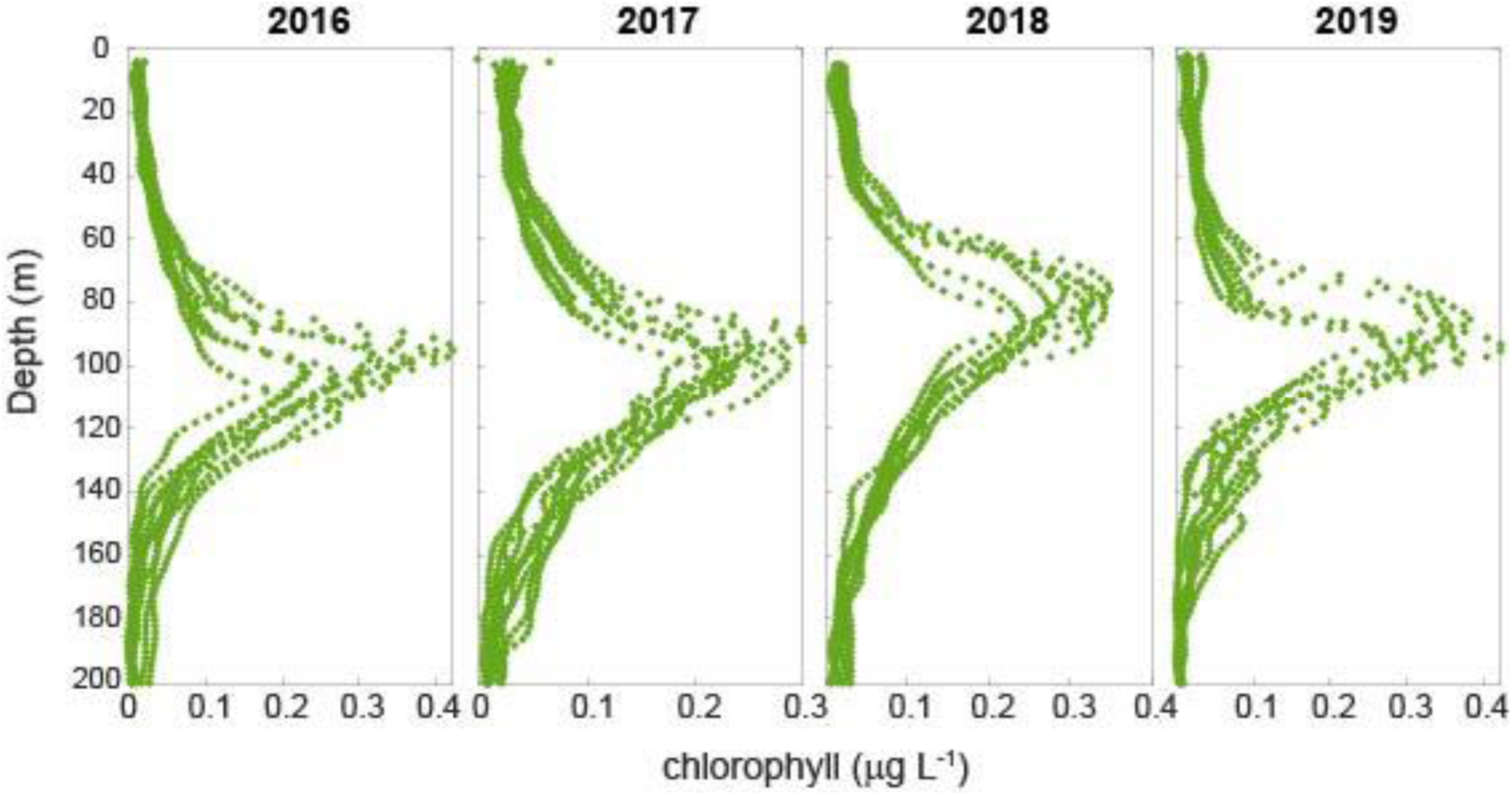
Chlorophyll fluorescence in the water column for each year, only showing the casts where metabolites were collected. Data have been processed into one-meter bins for each cast.

**Supplemental Figure 2.**
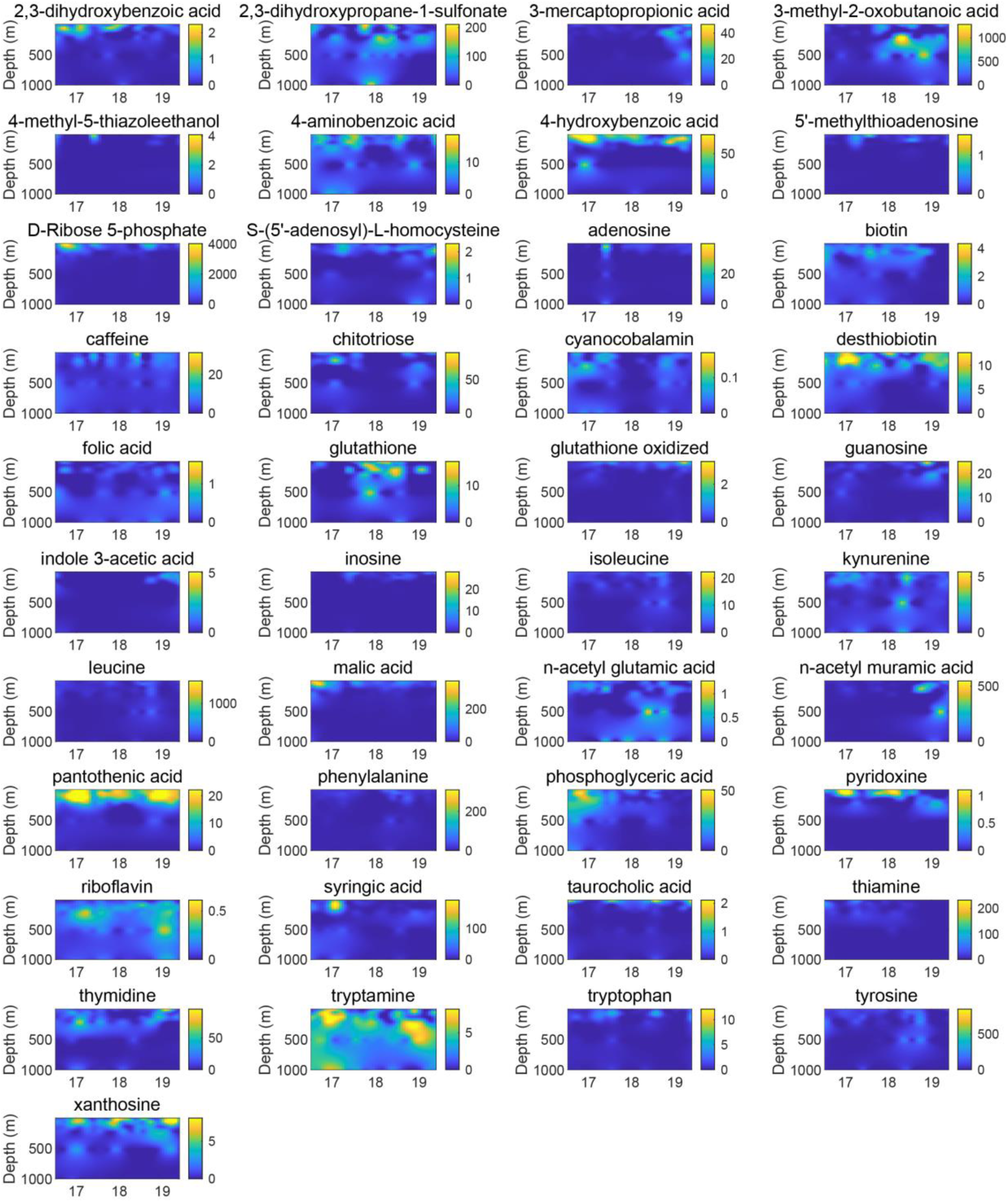
Color plots showing the complete set of metabolite data over all depths and years of the project. The color bars show the concentrations (in pM) for each metabolite.

**Supplemental Figure 3.**
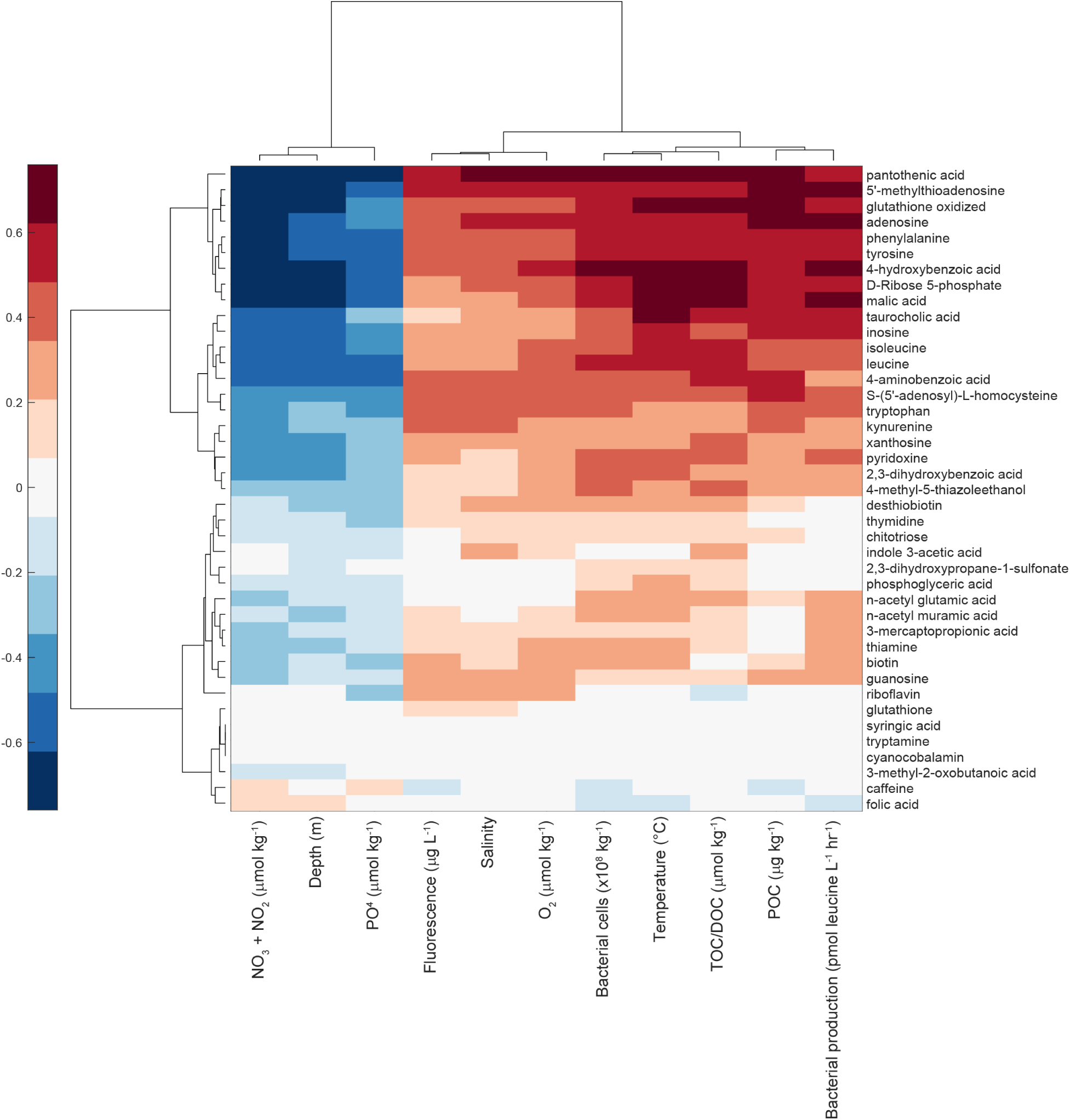
Correlations between select environmental parameters and the concentrations of metabolites measured over all four years of the project. Correlations are Spearman correlations, and the comparisons with *p*-values that are <0.05, after correction for a false discovery rate of 5%, are shown in the figure. Comparisons in white were non-significant.

**Supplemental Figure 4.**
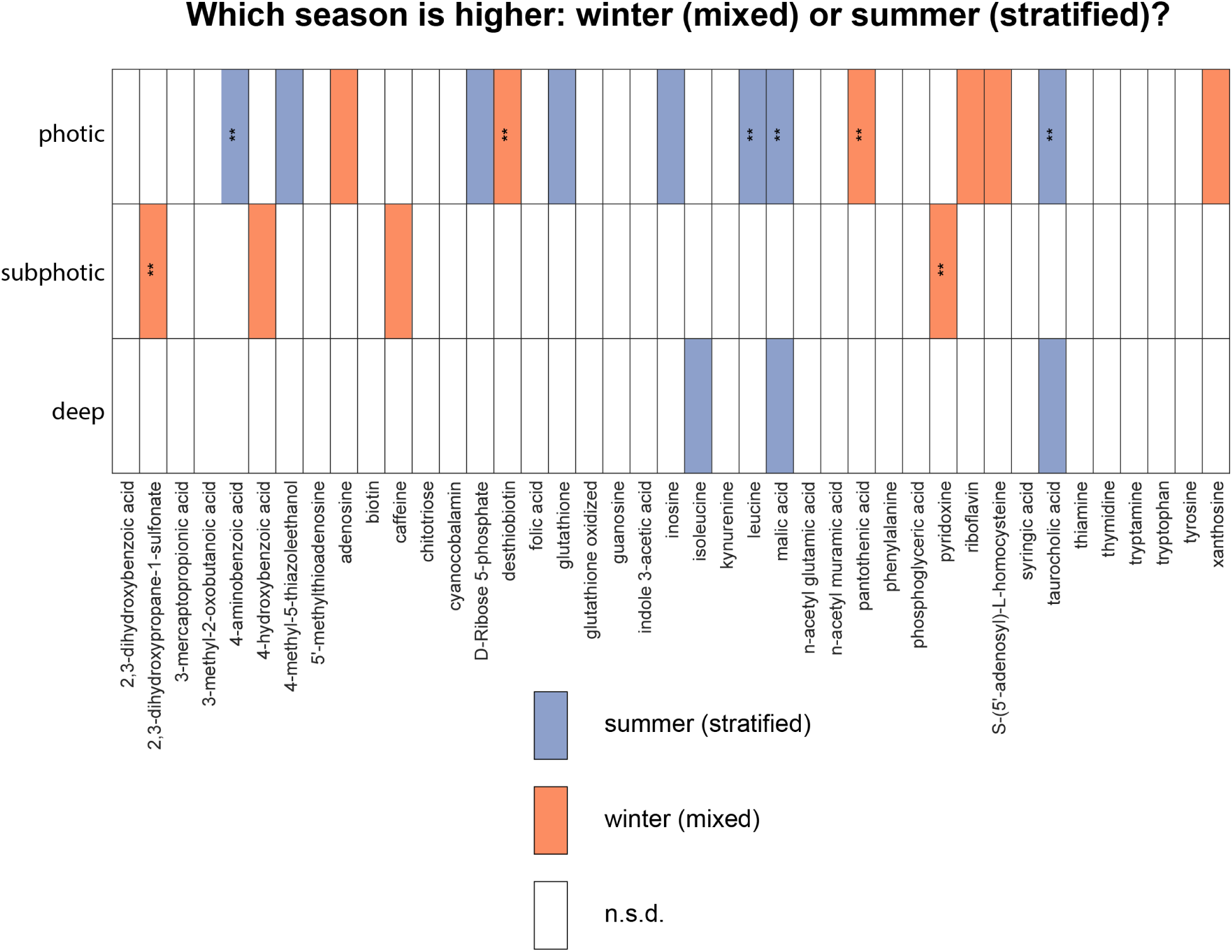
Summary differences between the winter (mixed) season and the summer (stratified) season in metabolites. Boxes in blue are dissolved metabolites where the summer season was higher, while orange shows the metabolites where the winter season was higher. In white are the cases where there was no significant difference between the two seasons. The water column was grouped into three regions for this comparison: photic zone (VZ_0_, VZ_1_, and VZ_2_), subphotic (VZ_3_), and deep (VZ_4_ through VZ_10_). Comparisons marked with ** have p-values <0.001, all other comparisons marked in orange or blue have 0.001 ≤ p-values < 0.05.

**Supplemental Figure 5.**
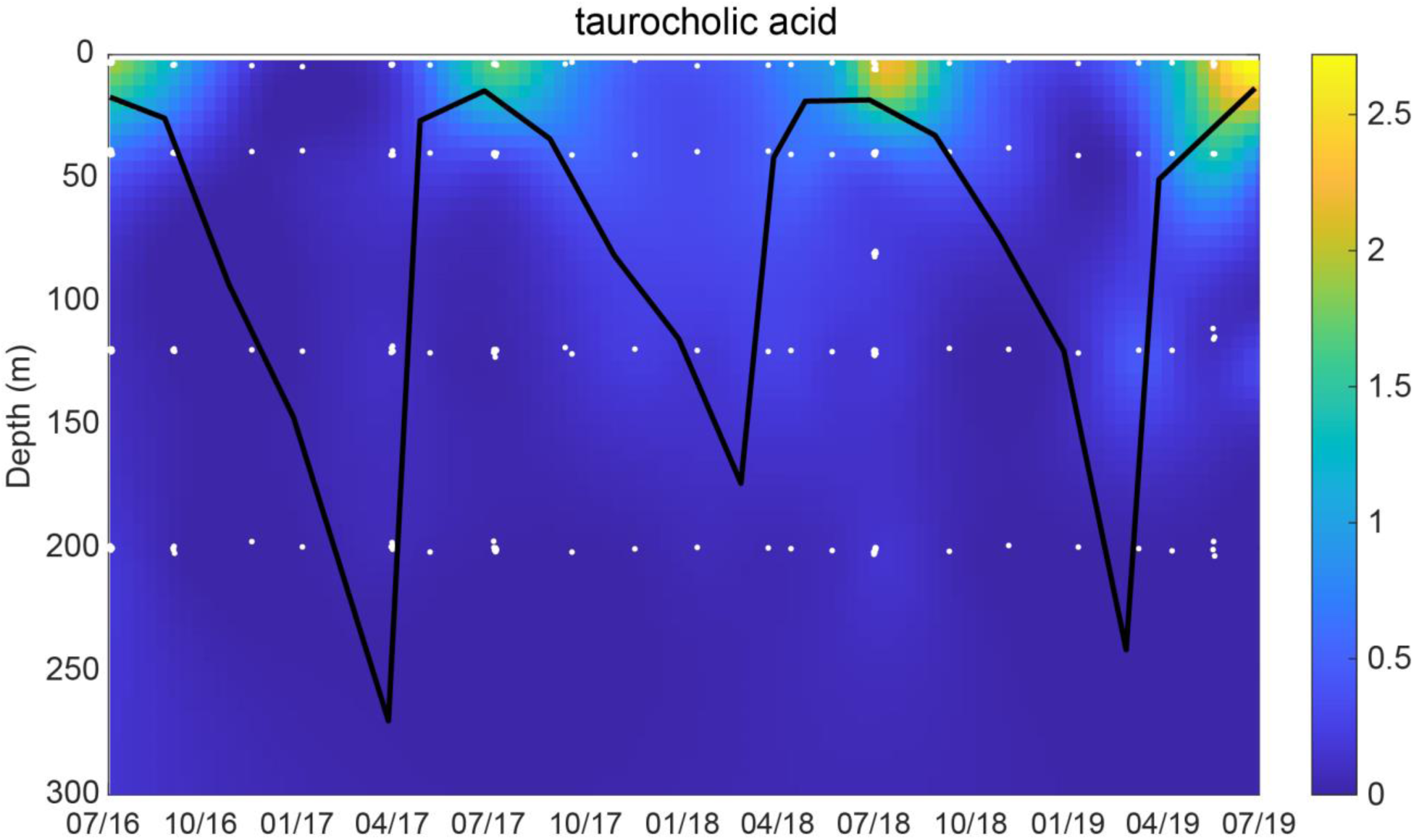
Dissolved taurocholic acid from July 2016 through July 2019 in the upper 300 m of the water column. The black line is the mixed layer depth. Colors represent concentration of taurocholic acid in pM.

**Supplemental Figure 6.**
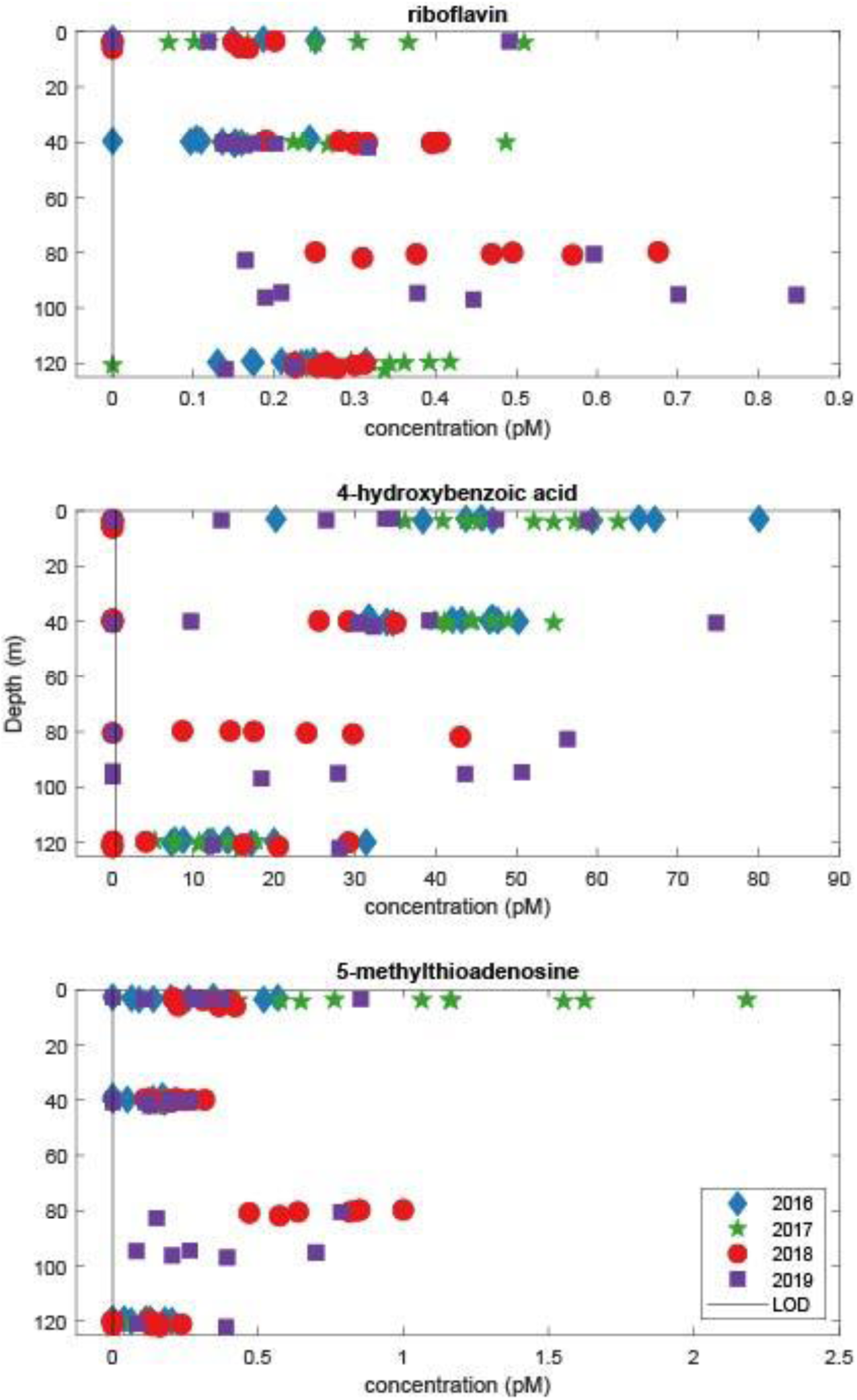
Concentrations (in pM) of riboflavin, 4-hydroxybenzoic acid, and 5-methylthioadenosine during the four July sampling periods. The vertical gray line is the limit of detection (LOD) for each metabolite. Additional sampling depths were added between 80 and 90 m in 2018 and 2019, and a different x-axis scale is used for each metabolite.

## Notes

### Competing Interest Statement

The authors have declared no competing interest.

